# Gamma band oscillations drives information flow in theta and delta band between hippocampus and medial pre-frontal cortex

**DOI:** 10.1101/809962

**Authors:** Samuel I.B.H.

## Abstract

Information flow in the brain is mediated by neuronal oscillations. Prior research has shown that gamma activity considered to be most reflective of neuronal firing is nested within theta cycles. This finding has given rise to the notion that theta phase mediates the gamma oscillations and thereby paces information flow. Such findings have been observed predominantly in cases where the gamma and theta oscillations were measured from the same underlying neuronal substrate. In this article, we analyze the across region, inter-brain coupling of gamma and theta oscillations using non-directional and directional connectivity measures. We show that in this context the information flow appears to be mediated by gamma oscillations which in turn drives the theta and delta oscillations. Additionally, different bands of gamma are coupled with theta and delta bands. During task, this information flow is enhanced compared to baseline in the frequencies which are coupled. Furthermore, the connectivity measures namely cross-frequency coupling and generalized partial directed coupling are not correlated with each other suggesting that they maybe representative of different underlying neuronal mechanisms.

## Introduction

Neuronal oscillations are intrinsically generated periodic patterns of neural activity. They are modulated by variations in task demand [1] and are involved in mediating information flow in the brain [2]. Prior research has shown prominent oscillatory activity in the delta (∼3Hz), theta (∼7Hz) and gamma (>30Hz) bands which have been associated with cognitive processes such as learning and memory [3]. Research in this field has led to the general notion that the high frequency gamma activity which reflects the ensemble firing of the neuronal substrates are paced by the low frequency theta activity [4]. Such findings led to the general notion that the information flow in the brain is mediated by the lower frequency oscillations. However, recent reports have challenged this notion by suggesting that the gamma oscillations drive the theta oscillations [5]. In this report we analyzed the interaction between the low frequency oscillations (theta and delta band) and the high frequency oscillations (gamma band) to characterize how they mediate information flow across different brain regions. The local field potentials from two brain regions namely, the hippocampus (dCA1) and the medial pre-frontal cortex (mPFC) was analyzed with connectivity measures such as the directed coherence (DC) and cross-frequency coupling (CFC) during baseline and task conditions.

### Theta and Gamma Oscillations in Hippocampus

The hippocampus is involved in memory related functions [6] and communicates with other regions to facilitate the learning process [3]. The predominant oscillatory activity in the hippocampus are the theta and gamma band oscillations [2]. The theta band activity is a slow oscillatory activity with a peak amplitude at about 7Hz and is present during active behavior [7]. Prior studies, show increased theta band activity during walking and during active participation in task with higher task demand resulting in higher theta activity [8]. The gamma band activity is a faster oscillatory neural activity which is more pronounced during high cognitive demand [9]. Since theta and gamma are dependent on task demand and their activity is correlated with each other. Therefore, these two oscillatory activities are believed to communicate with each other to execute the task [10]. These oscillatory activities play an active role in the processing of information within the hippocampus and in relaying information to other brain structures to facilitate goal-directed behavior.

### Delta and Gamma Oscillations in Medial Pre-Frontal Cortex

The medial Pre-Frontal Cortex (mPFC) is integral for executive functions such as decision making [11], goal-oriented behavior [12] and in selective attention [13]. The mPFC has direct projections from the hippocampus and works with the neural information coming from the hippocampus to perform its functions [14]. Previous studies show a dominant delta oscillation in mPFC which is modulated by task demand [15] and also by the activity in the hippocampus [16]. However, the exact mechanisms in which they communicate are not yet fully understood. Most studies use differences across task conditions to analyze the relationship between the mPFC and hippocampus which does not directly quantify the amount and direction of information flow.

### Neural Communication

Quantitative measures of neuronal interactions have been traditionally carried out using cross-correlation of the local field potentials (LFP). Recent work applied this approach across different frequency bands and the method is commonly referred to as cross-frequency coupling - CFC [17]. Studies using this method of analysis have shown that the amplitude of gamma activity is related to the phase of the theta activity [7], [18]. Furthermore, neuronal firing rate has also been shown to be dependent on theta phase [16]. Based on these studies, the general notion is that the theta phase drives the neuronal firing rate which reflects the gamma band power.

However, this method of analysis does not directly quantize the directional interaction. It is also not known how such interactions are affected by increases in task demand. Furthermore, prior research has largely focused on the interactions of the different frequency bands within an area. To understand the mechanisms of interaction of different frequency bands within a region and across regions, we performed CFC based analysis and compared it with a directional measure of connectivity namely: directed coherence (DC) on the LFPs from the dCA1 region of the hippocampus and the mPFC of mice [19]. Based on prior studies the following hypothesis was proposed and tested.

### Hypothesis

If theta activity influences the gamma activity, then it is expected that when CFC between these two oscillatory activities is high and the directional connectivity measure should predict a directional influence from the theta source to the gamma source. Furthermore, although these measures have been used widely in research, the relationship between them have not been characterized. If the two measures of connectivity are similar, then it is expected that the two measures namely, CFC and DC should be correlated.

## Materials and Methods

The data analyzed in this report was collected by the Peterson Lab [20] and a brief description of the experiment is provided in this report as follows. Chronic LFP recordings were obtained from mice during whisker stimulation at 6 sites namely, primary and secondary somatosensory cortex, primary motor area, parietal associative area, hippocampus (dCA1) and medial Pre-frontal Cortex (mPFC). Local field potentials (LFPs) from the dorsal CA1 and mPFC region (Figure 1A, 1B and 1C) were chosen for analysis since prior literature extensively characterizes oscillatory activity in these regions which helps interpret findings. All mice that had concurrent LFP data from both sites were considered for further analysis (n=13).

**Figure 1.**
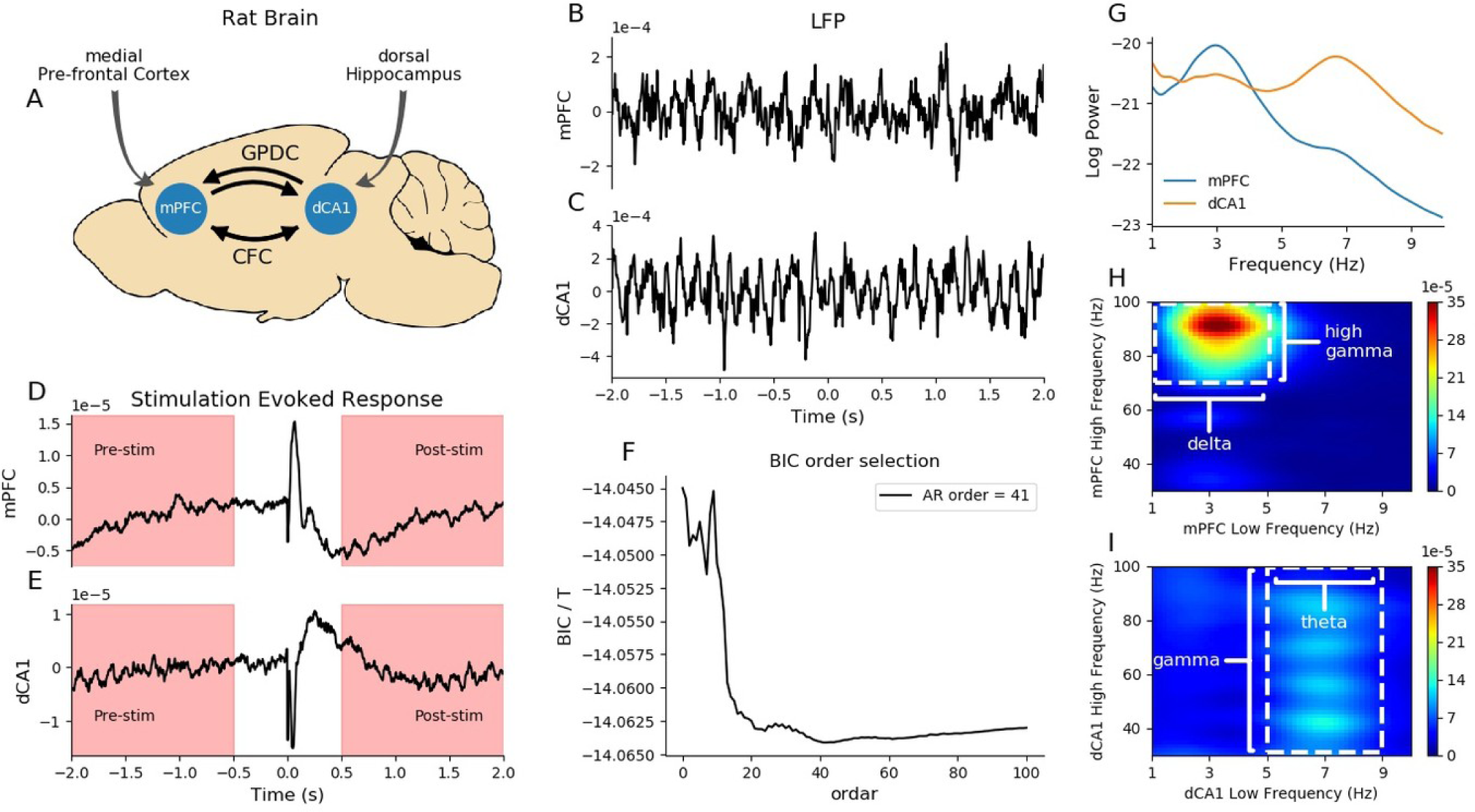
(A) Schematic of connectivity measures - Cross-Frequency Coupling (CFC) and Generalized Partial Directed Coherence (GPDC) and regions of Interest - medial Pre-Frontal Cortex (mPFC) and dorsal Hippocampus (dCA1) (B) Local Field Potential (LFP) from representative mPFC channel (C) LFP from representative dCA1 channel (D) Grand averaged Event Related Potential (ERP) from mPFC (E) Grand averaged ERP from dCA1 (F) Bayesian Information Criterion (BIC) as a function of AR model order (G) Power spectrum form both mPFC and dCA1 channels (H) Cross-frequency coupling (CFC) plot within mPFC and (I) CFC within dCA1

### Preprocessing

Each data sample had approximately one thousand two hundred whisker stimulation trials. The LFPs were then epoched to durations of 1.5s from before and after the stimulation (Figure 1D and 1E). To reduce line noise a notch filter at 50Hz was applied and then the signals were detrended. The epoch before stimulation is termed ‘pre-stim’ because of the lack of stimulus related processing. The epoch after the whisker stimulation started from 500ms post stimulus to remove the effects of the evoked potentials on the spectral measures. This epoch is termed ‘post-stim’ because of the preceding whisker stimulation. This period is associated with task related processing.

### Analysis

Power spectrum is obtained using the periodogram method and represents the magnitude of the oscillatory neural processes in the LFP recording. For each pair of electrodes cross-frequency coupling (CFC) measure is obtained using an auto-regressive (AR) model and the order of the AR model was estimated using the Bayesian Information Criterion (BIC) (Figure 1F) [21]. This measure is not a directional measure of oscillatory coupling. A directional measure of coupling namely - directed coherence (DC) was then obtained for inter-region electrode pairs [19] since DC cannot be calculated for signals from the same source.

### Non-Directional Connectivity Measure: Cross-Frequency Coupling

Cross-frequency coupling (CFC) is a measure of similarity across two frequency bands of activity. Since it is a bidirectional estimation, it does not provide any causal or directional connectivity estimate. Using this method, we obtain a CFC measure over a combination of different low frequency band and high frequency bands.

### Directional Connectivity Measure: Directed Coherence

Directed Coherence (DC) is a directional connectivity measure that quantifies the directional influence of the source signal on the sink signal. Since this is a directional measure between dCA1 and mPFC, we obtain two measures representing the directional influence - (i) dCA1 -> mPFC and (ii) mPFC -> dCA1.

These two measures were then correlated with each other for pre-stim and post-stim epochs to test whether they reflect the same underlying mechanism using Pearson’s correlation. For these correlations frequency range of interest were defined as follows based on the observed CFCs: (i) delta: 1Hz-5Hz, (ii) theta: 5Hz-9Hz, (iii) gamma: 30Hz-100Hz, (iv) high-gamma: 70Hz-100Hz.

## Results and discussion

### Pre-stim Period

#### Power spectrum and within region CFC

During pre-stim time period, we see that dCA1 has a prominent theta band activity while the mPFC has a prominent delta band activity (Figure 1A) which is in agreement with prior literature [18]. First within region oscillatory interactions were analyzed. In this case, DC is not applicable because it requires two signals having different sources to estimate directional influence. However, it is possible to obtain the CFC since it can quantify connectivity at different frequencies within the same signal.

Within mPFC, the CFC analysis shows that the gamma band activity is correlated with delta phase (Figure 1B). Similarly, within dCA1, gamma band activity is correlated with the theta band phase (Figure 1C). This is in agreement with prior literature which shows that the local low frequency oscillations are coupled with the local high frequency oscillations [17]. However, there is a difference in the gamma band frequencies that are associated with the lower frequency: (i) In mPFC, high gamma activity (70Hz-100Hz) is associated with the low frequency activity whereas (ii) in dCA1, gamma activity over a broad range (30Hz-100Hz) is associated with low frequency activity. Additionally, the CFC within mPFC (delta - high gamma; 2B) is much larger (2J) than the CFC in dCA1 (theta - gamma; 2C).

#### Inter-region connectivity

For cross-region interaction we compare CFC and DC between dCA1 and mPFC to characterize their relationship. The conventional notion is that the mPFC theta is modulating the gamma power in dCA1 since lower frequency oscillations are thought to modulate the higher frequency oscillations [7], [12], [16], [18]. This implies that the causal influence is from mPFC to dCA1 in the theta band. Contrary to this notion, we observe the following: (1) Figure 2D - DC shows that the causal influence is from dCA1 to mPFC in the theta band and (2) Figure 2E – CFC shows that dCA1 gamma is coupled with mPFC theta. Together, these findings suggest that the increased coupling (CFC) between dCA1 gamma and mPFC theta may be due to the causal influence of dCA1 gamma on mPFC theta. Similarly, CFC is pronounced between mPFC gamma and dCA1 delta and the DC from mPFC to dCA1 is pronounced at the delta band. This is also in line with the notion that the higher frequencies (mPFC gamma) influences lower frequencies (dCA1 delta). For within region coupling, CFC between mPFC delta and mPFC high gamma is greater than dCA1 theta and dCA1 gamma (Figure 2G) whereas for cross-regional coupling the relationship is inverted (Figure 2H). Additionally, the within region CFC seem to be always greater than cross region CFC (Figure 2I and 2J).

**Figure 2.**
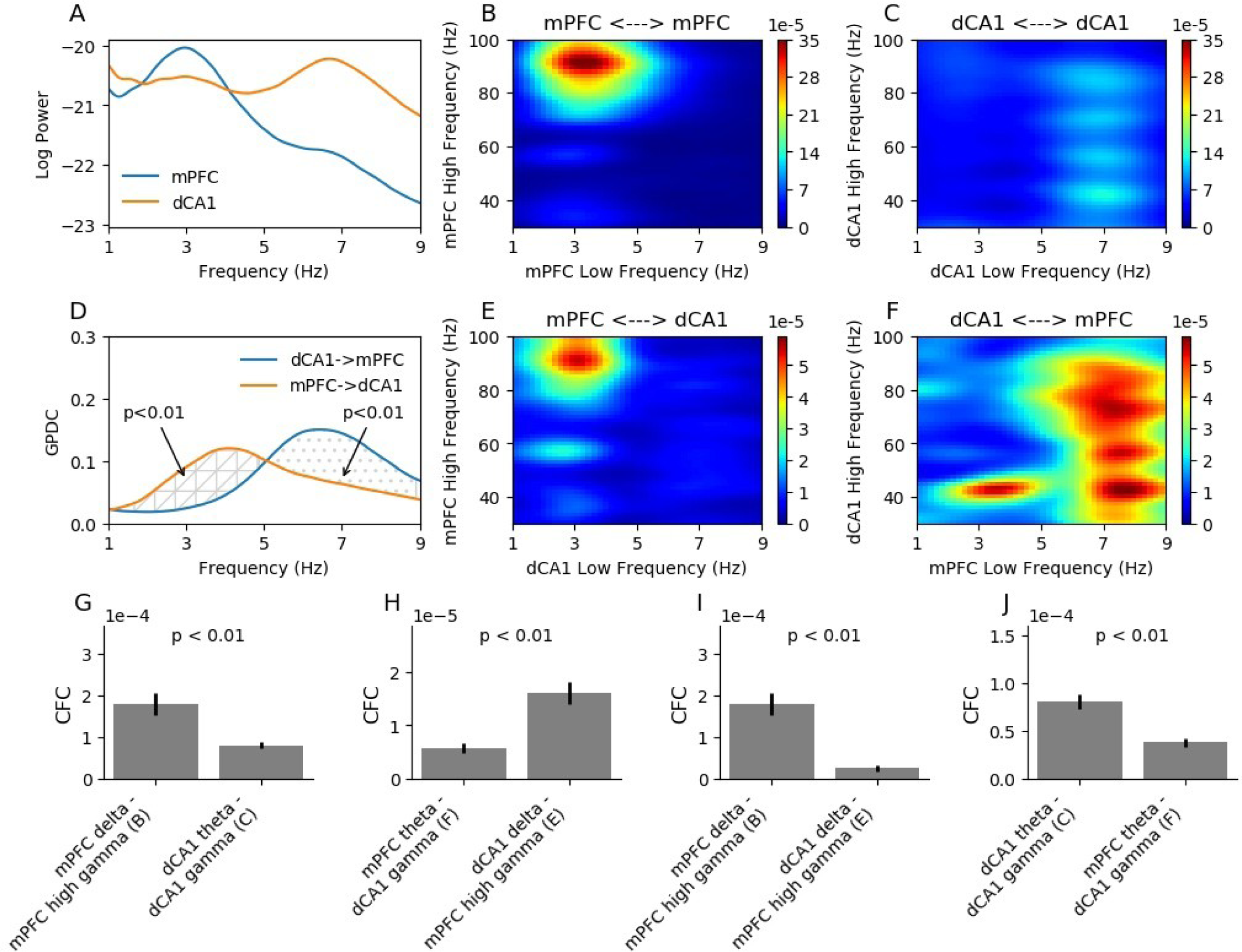
Oscillatory interactions between mPFC and dCA1 before stimulation: (A) Power spectrum of dCA1 and mPFC. (B) CFC within mPFC (C) CFC within dCA1. (D) DC from dCA1 to mPFC. (E) CFC between mPFC high frequency power to dCA1 low frequency phase. (F) CFC from dCA1 high frequency power to mPFC low frequency phase. (G) Within region CFC difference (B vs C). (H) Cross region CFC difference (E vs F). (I, J) Difference between within region and cross region CFC.

#### Relationship between CFC and DC

To determine whether the CFC and the DC are measuring the same underlying mechanisms, the two measures were correlated. However, no significant correlations were found between the DC at theta from dCA1 to mPFC and the CFC between dCA1 gamma and mPFC theta (r=.01, p=.97). Similarly, the DC at delta from mPFC to dCA1 was not correlated with CFC from mPFC high-gamma to dCA1 delta (r=.29, p=.34). This may be due to the CFC being more focused in a smaller frequency band from mPFC high-gamma to dCA1 delta, whereas for dCA1 gamma to dCA1 theta the CFC is pronounced over a wider gamma frequency.

#### Post-stim Period

The same analysis as described for the pre-stim epoch was carried out for the post-stim period and similar findings were observed which suggests that the gamma source influences the theta/delta source (Figure 3 A-F). These results suggest that the patterns of oscillatory interactions are similar across pre-stim and task related cognitive states. Furthermore, no significant correlations were found between the DC at theta from dCA1 to mPFC and the CFC between dCA1 gamma and mPFC theta (r=.05, p=.88). Similarly, the DC at delta from mPFC to dCA1 was not correlated with CFC from mPFC high-gamma to dCA1 delta (r=.01, p=.99).The amplitude differences between the CFC measures during post-stim period (Figure 3G-J) also seem to follow the same pattern as seen in the pre-stim period (Figure 2G-J).

**Figure 3.**
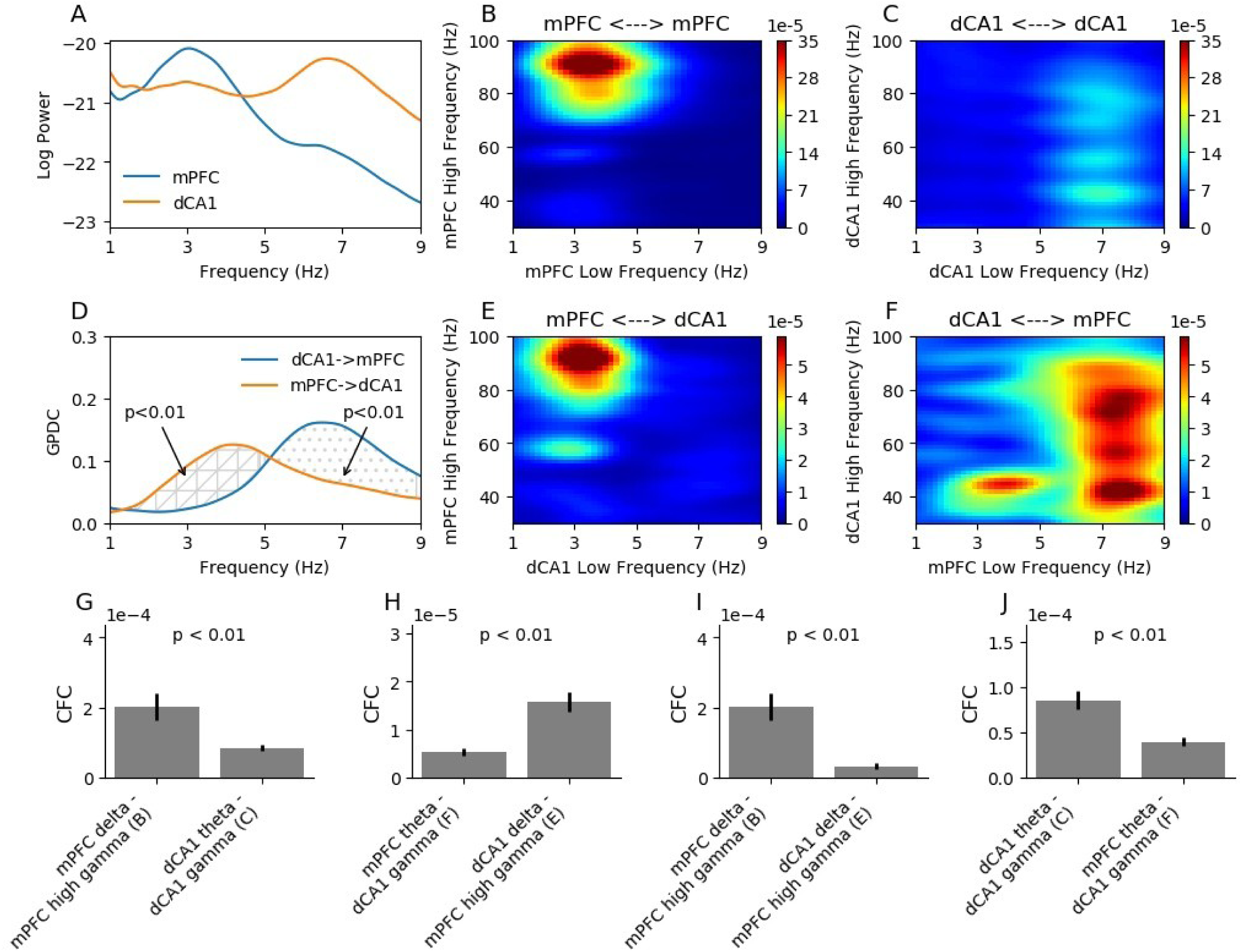
Oscillatory interactions between mPFC and dCA1 after stimulation: (A) Power spectrum of dCA1 and mPFC. (B) CFC within mPFC (C) CFC within dCA1. (D) DC from dCA1 to mPFC. (E) CFC between mPFC high frequency power to dCA1 low frequency phase. (F) CFC from dCA1 high frequency power to mPFC low frequency phase. (G) Within region CFC difference (B vs C). (H) Cross region CFC difference (E vs F). (I, J) Difference between within region and cross region CFC.

### Comparison of Post-stim period to pre-stim Period

#### Power spectral changes

Since whisker stimulation evokes sensory and associated cognitive processes, it is expected that the communication between dCA1 and mPFC are enhanced due to the higher cognitive demand. To test this notion, the differences in the connectivity measures between pre-stim and post-stim periods were analyzed. The results show that in mPFC there is an overall increase in the high-delta band power, whereas the low-delta band power decreased (Figure 4A). Therefore, in the following analysis to reflect this increase in the delta band peak frequency, a high-delta band (3-5Hz) was defined. In dCA1, during post-stim compared to pre-stim, there is overall decrease in the lower band power, with the theta band power decreasing the least (Figure 4A). This result suggests that the brain is able to selectively enhance or suppress certain oscillatory activity to meet the cognitive demand. Furthermore, there is selective enhancement of specific frequencies even within the classically identified oscillatory activity. Though the dominant delta band peaks at about 3Hz, estimates of causal interaction suggests that information flow is enhanced in the high delta band with a peak at about 4Hz (Figure 4G). The differential increase in high-delta compared to low-delta power may be due to (1) multiple generators of the delta band activity or (2) the task induced increase in neurotransmitters and processing speed which is known to increase the frequency of oscillatory activity in the brain [22].

**Figure 4.**
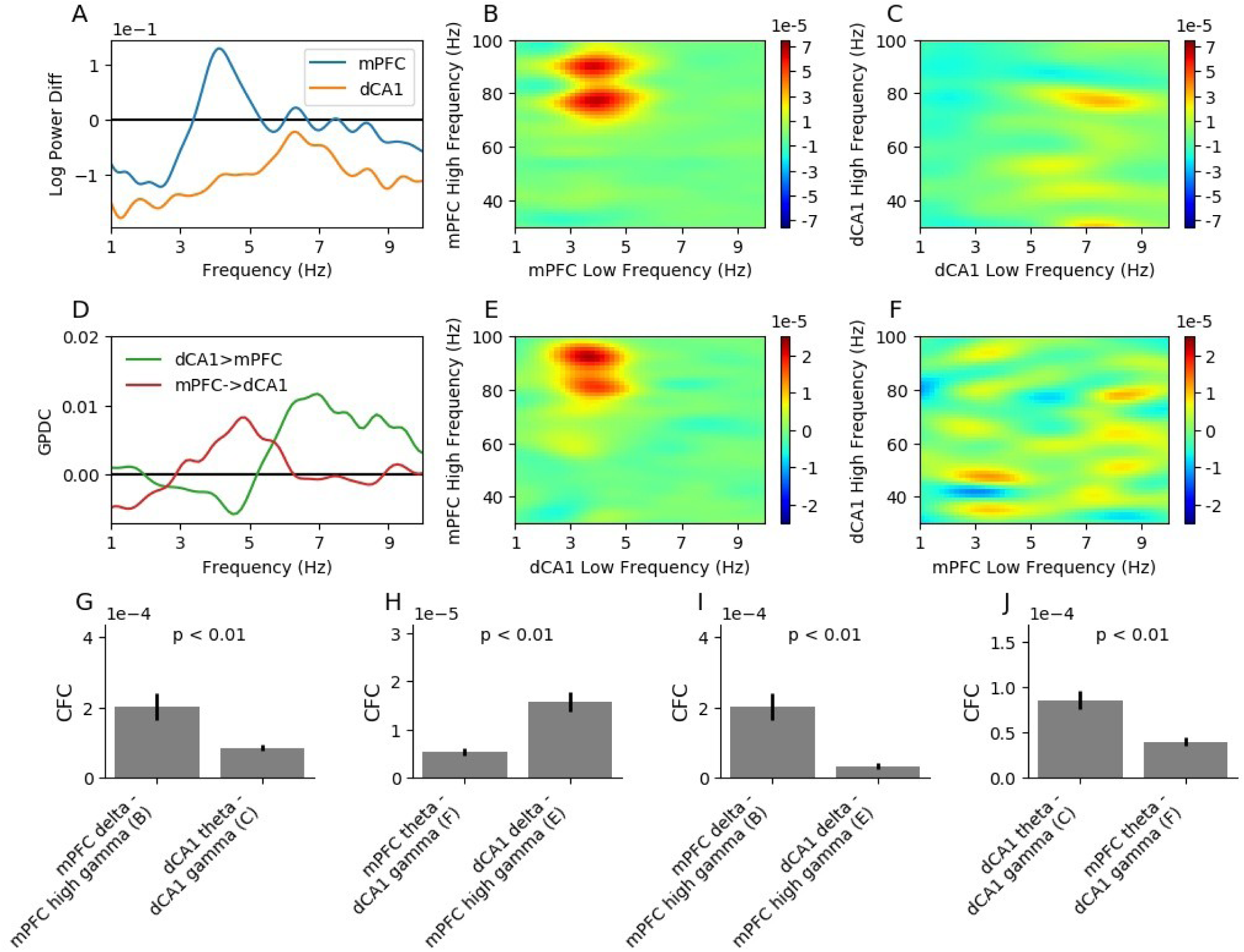
Differences in oscillatory interactions between mPFC and dCA1, pre and post stimulation: (A) Power spectrum difference in dCA1 and mPFC. (B) CFC difference within mPFC (C) CFC within dCA1. (D) DC difference from dCA1 to mPFC. (E) CFC difference between mPFC high frequency power to dCA1 low frequency phase. (F) CFC difference from dCA1 high frequency power to mPFC low frequency phase. (G) Within region CFC difference (B vs C). (H) Cross region CFC difference (E vs F). (I, J) Difference between within region and cross region CFC.

#### Within region CFC

Within region CFC reflects the change in oscillatory power at both mPFC and dCA1. In mPFC, there is an increase in CFC in the high-delta band similar to the power change (Figure 4B). The CFC suggests that the increase in high-delta is associated with high-gamma band frequency and thus provides additional information not evident in the power spectrum. In dCA1 changes in CFC (Figure 4B) are not as prominent as that seen in mPFC (Figure 4B).

#### Cross-region connectivity

Theta band DC from dCA1 to mPFC increases in the post-stim period compared to pre-stim (Figure 4D). The CFC between dCA1 gamma and mPFC theta also shows increase (Figure 4E). However, the CFC and DC measures at theta band are not correlated (r=.09, p=.78). Similarly, DC from mPFC to dCA1 increases in the post-stim period compared to pre-stim in the high-delta band (Figure 4D). There exists a consistent increase in CFC from mPFC high gamma to dCA1 high-delta (Figure 4E) but the CFC and DC measures are not correlated (r=.24, p=.44). These results suggest that the CFC and DC are different underlying measures of connectivity and taken together is able to provide a better characterization of inter-region brain connectivity. The amplitude differences between the CFC measures between post-stim and pre-stim period (Figure 4G-J) also seem to follow the same pattern as seen in the pre-stim (Figure 2G-J) and post-stim periods (Figure 3G-J).

#### Increased CFC between gamma and theta is associated with directional influence from gamma to theta

The results presented in this report confirms previous findings that there is increased inter-region connectivity between high frequency activity and lower frequency activity (delta, theta). However, the directional connectivity results suggest that the information flow occurs from the high frequency activity to the lower frequency activity which is contrary to the classical notion that the lower frequency activity drives the higher frequency activity. Further characterization of the inter-region connectivity and development of new techniques to measure directional connectivity across frequency domains are necessary to understand the modes of communication within and between cortical regions.

Furthermore, although CFC and the DC are related in certain cases, comparison across task and pre-stim shows that the two measures do not reflect the same underlying mechanisms. The results presented here suggests that directional connectivity measures are required to infer information flow even if two regions have high CFC between then at certain frequencies. Similarly, even if information flow can be obtained from directional connectivity measures, the exact frequencies at which such information takes place is better understood with CFC measure. For example, while DC show increased information flow after simulation compared to pre-stim, the CFC shows the frequencies at which this information flow might occur.

## Limitations

The low sample size is a limitation of this study and the lack of correlation between the DC and CFC maybe due to the lower sample size. However, even at low sample sizes we know that these are not highly related due to low r-values and consequently we suggest that they reflect different underlying processes. Post stimulation there are changes in the power distribution which changes the definition of frequency bands. For example, the low-delta is suppressed whereas the high-delta is enhanced after stimulation. These changes while consistent within the data, do not lend itself to a simple definition of oscillatory rhythms and consequently needs detailed inspection to interpret findings.

## Summary

The findings in this report suggests that contrary to the classical notion, it is the gamma band activity that drives the theta and delta band activity across brain regions. The CFC and DC measures are not correlated and therefore reflects different underlying neural mechanisms. While the cross-frequency coupling (CFC) seems to reflect the frequencies at which inter-cortical communications takes place, the directional coherence (DC) measures the direction and amount of information flow. The results provide evidence towards the higher frequencies driving the activity of the lower frequencies. Furthermore, the different measures of cross-region connectivity are not correlated and seem to reflect distinct underlying properties.

## Disclaimer

The views expressed are those of the author and do not reflect the official views of the University of Florida, Henry M. Jackson Foundation for the Advancement of Military Medicine, Inc., the War Related Illness and Injury Study Center or the Veteran Affairs Medical Center, Washington, DC.

